# MathIOmica: an integrative platform for dynamic omics

**DOI:** 10.1101/074260

**Authors:** George I. Mias, Tahir Yusufaly, Raeuf Roushangar, Lavida R. K. Brooks, Vikas V. Singh, Christina Christou

## Abstract

Multiple omics data are rapidly becoming available, necessitating the use of new methods to integrate different technologies and interpret the results arising from multimodal assaying. The MathIOmica package for *Mathematica* provides one of the first extensive introductions to the use of the Wolfram Language to tackle such problems in bioinformatics. The package particularly addresses the necessity to integrate multiple omics information arising from dynamic profiling in a personalized medicine approach. It provides multiple tools to facilitate bioinformatics analysis, including importing data, annotating datasets, tracking missing values, normalizing data, clustering and visualizing the classification of data, carrying out annotation and enumeration of ontology memberships and pathway analysis. We anticipate MathIOmica to not only help in the creation of new bioinformatics tools, but also in promoting interdisciplinary investigations, particularly from researchers in mathematical, physical science and engineering fields transitioning into genomics, bioinformatics and omics data integration.

## Introduction

With the advent of readily available omics technologies that will greatly aid the advancement of the emerging field of precision medicine, the need to integrate information from these disparate omics technologies^1–6^ (genomics, transcriptomics, proteomics, metabolomics, etc.) is becoming more apparent. The role of bioinformatics in analyzing such high throughput omics data is unquestionable, and evident in major projects, such as the ENCODE Project^7^, 1000 Genomes^8^, UK10K^9^ project, and will be indispensable in the Precision Medicine Initiative^10^ (PMI) now underway. In particular, novel medical insights are expected through the integration of genomic information with the global monitoring of molecular components and physiological states in a coherent fashion, and the modeling of the integrated complex systems and associated dynamic pathways. The molecular component interactions are central in all biological processes and have contributed to our rudimentary understanding of disease onset, progression and treatment. Large-scale efforts to globally follow all such omics components in systems and in individuals are currently underway, including genomic and pharmacogenomic considerations^11–16^. One of the first such examples was the integrative Personal Omics Project (iPOP)^16–18^, profiling multiple omics datasets from a single individual over multiple time points. These studies provide a plethora of data and more are on the way. However, the studies show that the methodology for integrating such information is underdeveloped, especially in terms of dynamical analyses that directly address the complexities of biological experimentation, such as uniform normalization procedures across time course omics data and uneven time sampling.

Great progress has been made in the area of development of bioinformatics tools and platforms, towards data integration^19–21^. Notable examples include Bioconductor^22, 23^ and BioPython^24^, Galaxy^25^, GenePattern^26^, DAVID^27^, QIAGEN’s Ingenuity Pathways (IPA®, QIAGEN Redwood City, www.qiagen.com/ingenuity), Cytoscape^28^ and many more. Bioinformatics is now an essential tool for the modern geneticist, and intertwined with every aspect of genomics research. The practicing bioinformatician typically uses a combination of programs for the job at hand, and their language of choice for development includes high-level languages such as R^29^ and Matlab by Mathworks, scripting languages such as Python and Perl, Unix shell scripts (e.g. Bash), as well as coding tools written in C, C++, Java, etc. Bioinformatics tools are continuously being developed and improved to address the increasing demand for sophisticated analysis tools. However there has not been as much development for bioinformatics in the Wolfram Language and Mathematica^30^. Mathematica, which was released in 1988^31^, has been widely used by mathematical and physical scientists and students and has extensive symbolic, statistical and computational capabilities. Furthermore, it is used widely in introductory and advanced mathematics courses, including first college courses in calculus. There have been a few packages for bioinformatics in Mathematica^32–36^, but a general approach and package, incorporating standard tools used in data exploration by biologists, has not been implemented systematically. We believe that the availability of bioinformatics packages in Mathematica will not only provide additional tools for current bioinformatics users, but additionally encourage interdisciplinary research from mathematical and physical science investigators that want to enter the field of applied bioinformatics.

In this work we present MathIOmica, an open-source software package written in the Wolfram Language for Mathematica that provides a framework for the analysis and interpretation of (dynamic) multi-omics data. The package is one of the first steps towards the development of new universal methods and tools to integrate biological omics data in the Wolfram language. It supplements Mathematica with multiple new functions that facilitate the analysis and development of new analysis methods for dynamical omics data. MathIOmica provides a framework for importing datasets in a structured way, including methods for processing such data, addresses time series analysis of omics data (including considerations for missing values), and includes annotation capabilities using known databases (such as UCSC Browser Tables^37, 38^, Gene Ontology^39, 40^ and KEGG pathways^41, 42^). A particular feature of MathIOmica is the extensive documentation with over 1000 pages total of tutorials and input/output examples for functions, including the options for each function, that have been inbuilt into the package and are readily available through Mathematica’s native help system upon installation. Two fully worked examples are also in the documentation based on dynamic data from the pilot iPOP project to provide additional real data examples of using the framework. MathIOmica was created with an outlook to become an extensible framework, to enable bioinformatics development in the Wolfram language, and use the considerable computational capabilities already available in Mathematica.

MathIOmica is available to download at https://github.com/gmiaslab/mathiomica and has a dedicated page found at https://mathiomica.org.

## MathIOmica’s Framework

### Overview and Workflow

We have been developing an integrative framework, MathIOmica, with multiple modules for omics downstream statistical analysis now completed. MathIOmica has multiple functions, Figure 1, utilizes a flexible data format, Figure 2, can implement multi-omics analyses, as shown in the example workflow in Figure 3, and provides various graphical interfaces and result visualizations, Figure 4. MathIOmica integrates multiple omics information starting from mapped experimental omics data - typically RNA-Sequencing expression levels, mapped protein intensities, and small molecules intensities. Using this framework we can analyze different omics data (genome, transcriptome and proteome) individually, based on each technology’s requirements, perform quality control (accounting for experimental and technical limitations) and set all the different technologies on common ground (statistical transformations). MathIOmica provides classification methods to identify patterns in the data, as well as annotation capabilities as discussed briefly below. Finally, extensive documentation is provided for every function and its option set.

**Figure 1.**
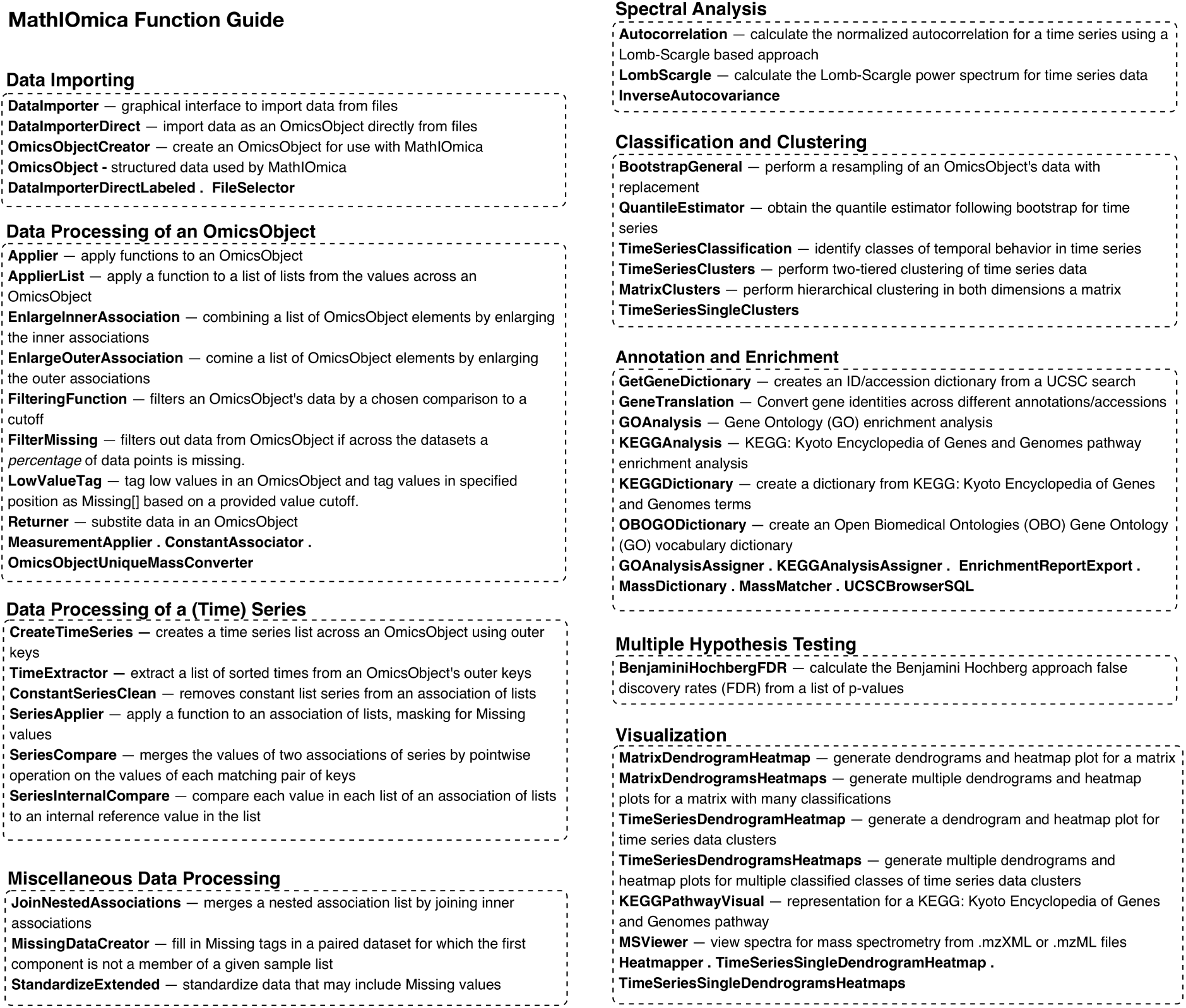
MathIOmica Guide. MathIOmica introduces multiple new functions to assist with bioinformatic analysis. The brief guide provides a short description for main functionality, and is available in the MathIOmica in-built documentation, providing a version with links pointing to each function’s detailed description and example usage. MathIOmica’s full documentation is integrated with Mathematica’s native documentation upon installation and can be evaluated in place by the user.

**Figure 2.**
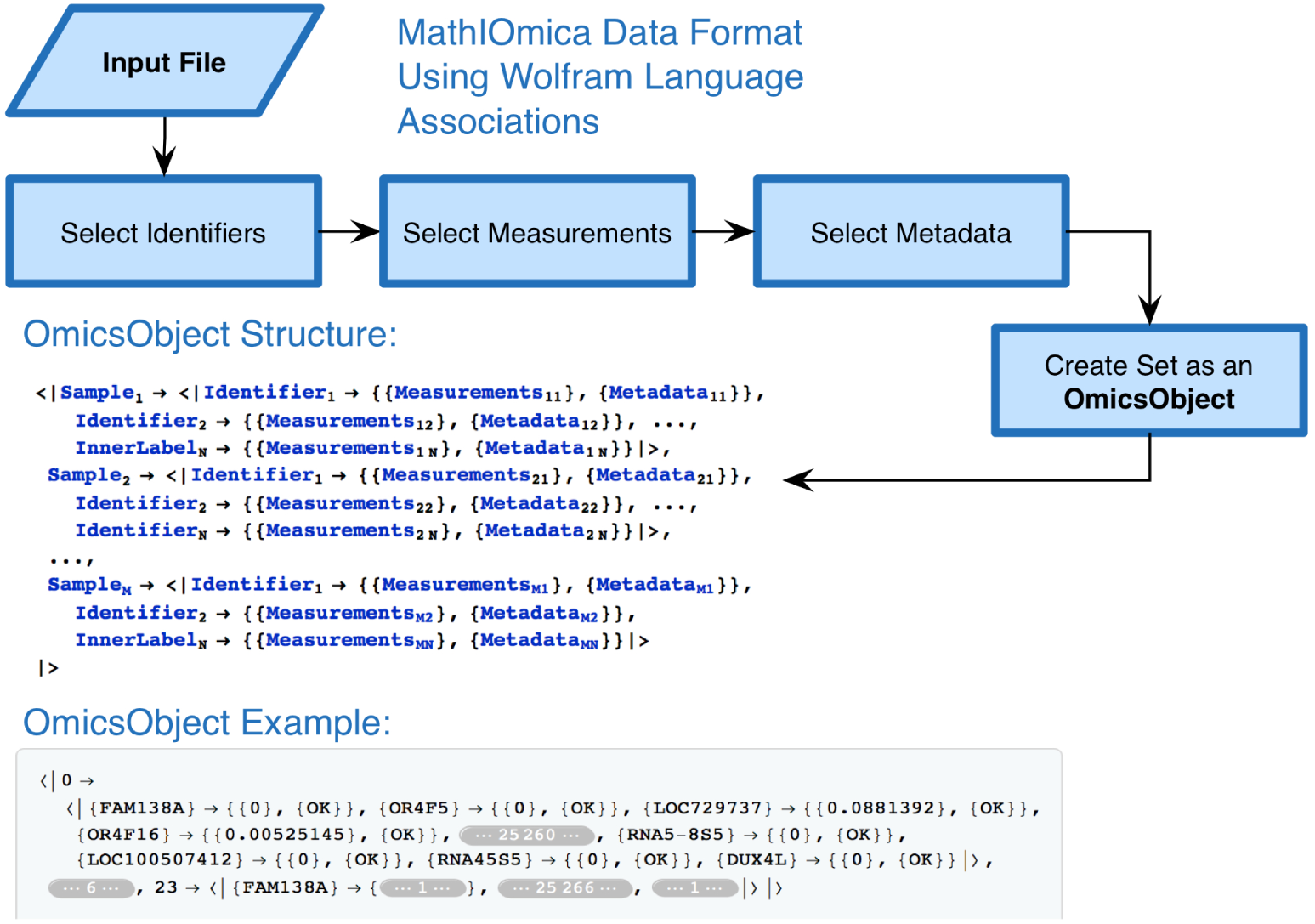
MathIOmica OmicsObject Format. The MathIOmica v.1 input format was redesigned to implement the 10.3+ Wolfram Language improvements to use associations (cf. dictionaries in Python). The data format is termed an OmicsObject, and is an association of associations with common identifiers as inner association keys across multiple samples. It addresses rapid sample identification, measurement and metadata all maintained for user accessibility and additionally accommodates missing values. In addition to the user having control of all aspects of the input data via Query commands on the OmicsObject representing the data, MathIOmica includes specialized functions designed specifically for creating and manipulating an OmicsObject.

**Figure 3.**
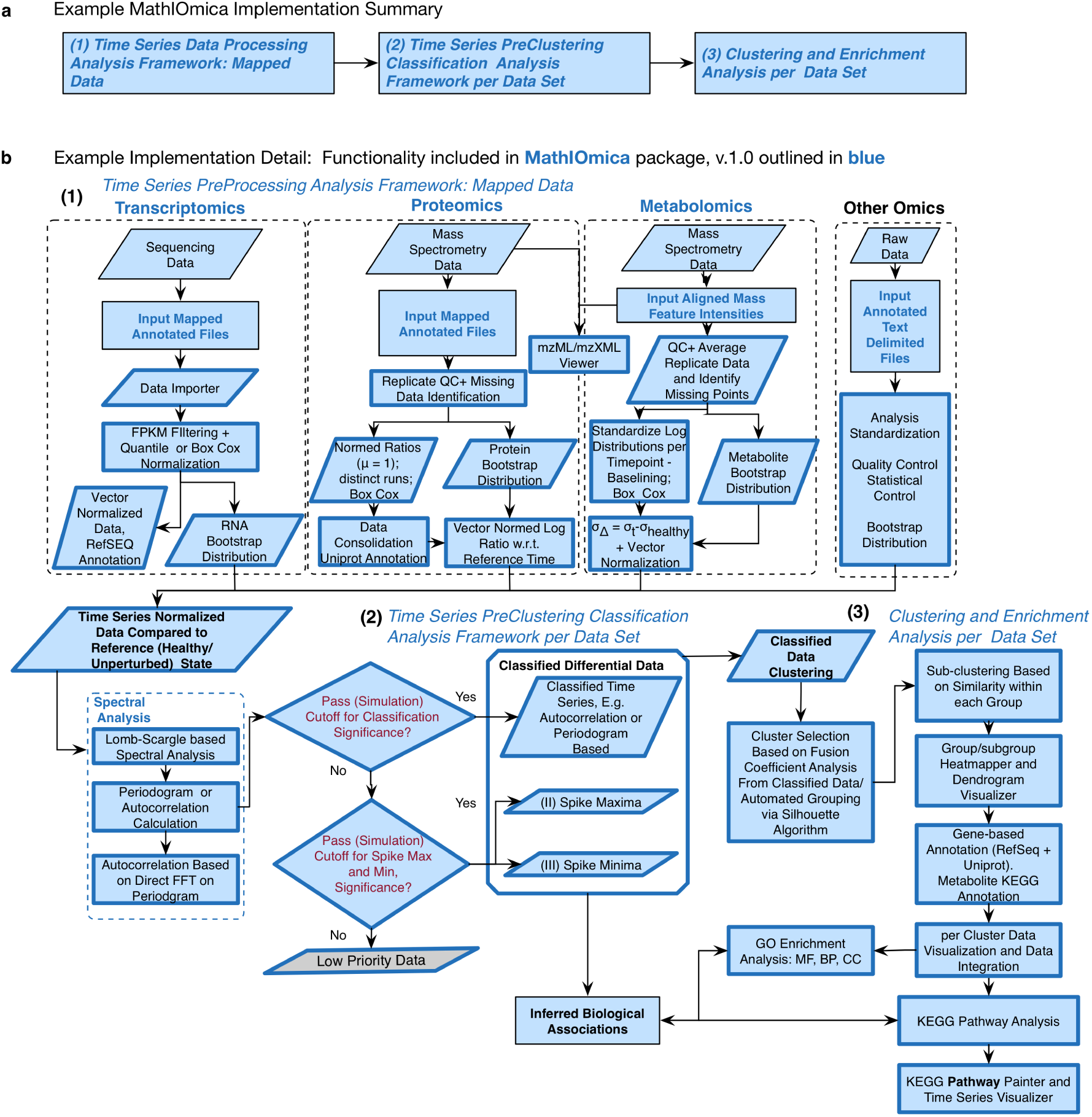
MathIOmica Implementation Example. MathIOmica allows the integration of multiple omics dynamic data. The generalized approach summarized in **a** first preprocesses each omics dataset according to its own considerations towards a common format of a time series. The time series can then be classified for temporal patterns using spectral analyses. Finally classes of temporal patterns are clustered, and the results can be visualized and further analyzed for Gene Ontology or KEGG pathway overrepresentation. A fully worked example with the various details shown in **b** is provided in the MathIOmica Tutorial as part of the in-built package documentation (see also Supplementary Note 1 for a printout version).

**Figure 4.**
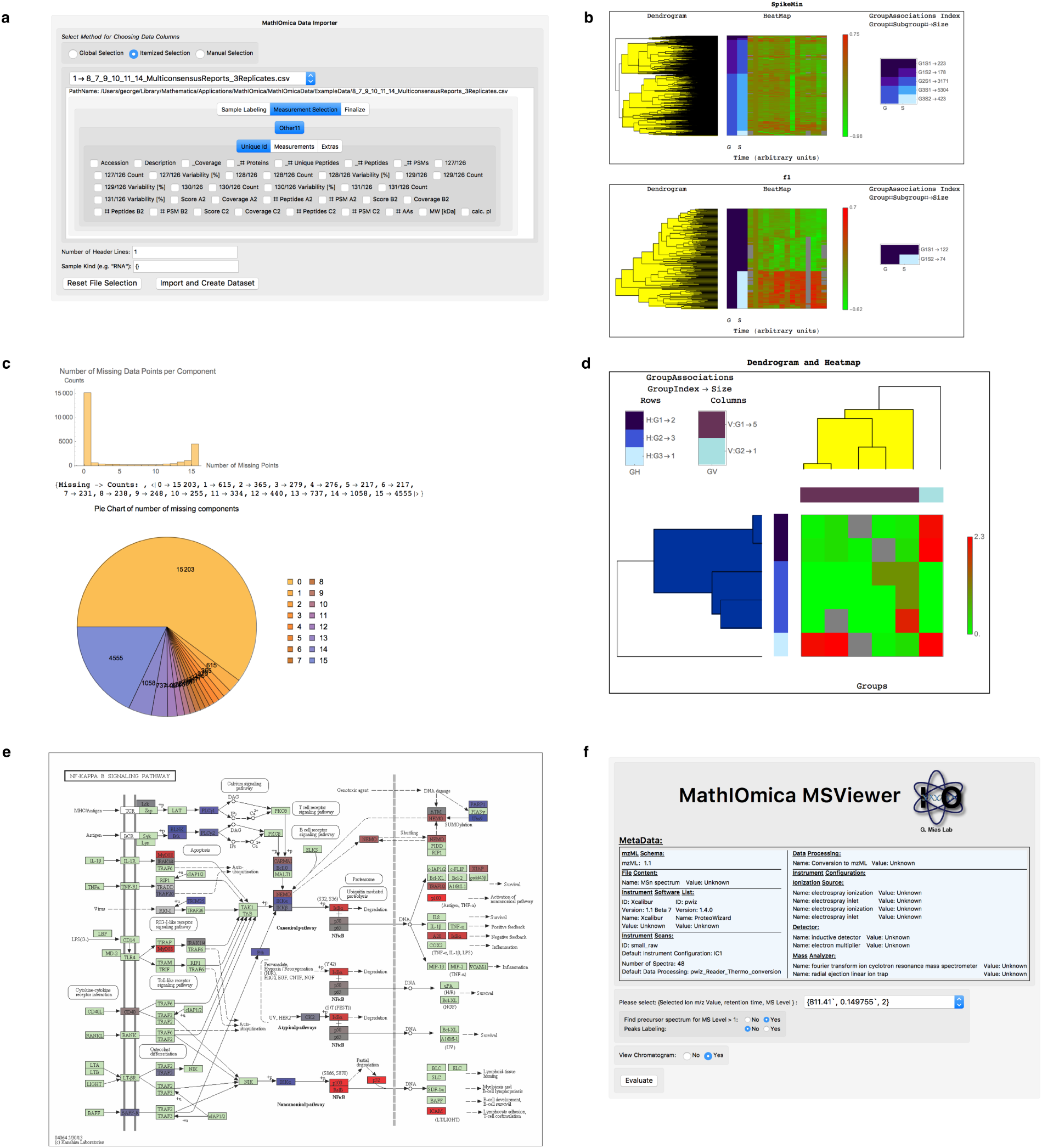
MathIOmica Example Screenshots. MathIOmica has various graphical interfaces and visualization capabilities, including: **a**, a graphical importer, **b**, time series clustering visualization, **c**, missing data filtering summaries, **d**, clustering of matrices, **e**, visualization of KEGG pathways, and **f**, a mass spectrometry viewer.

### Wolfram Language Code Base

MathIOmica was written exclusively in the Wolfram Language^31^. The language provides a robust, fully tested, cross-platform environment (Mac OS, Windows and Linux have been tested). The Wolfram Language already leverages symbolic, statistical, computational and database capabilities that are utilized by MathIOmica. The functions were written using the recently available association constructs (akin to dictionaries in other languages, e.g. Python), in Mathematica 10.4+^30^, and with a functional approach in mind. The source code is provided with the package, and adheres to standard Wolfram Language conventions with respect to capitalization and definitions of functions. All documentation and data files necessary are also provided with the code. In implementing the package in Mathematica, we find the standard Mathematica notebook interface provides a balance of detailed note taking, and rationale interweaved with code, commenting and results, and believe this feature to be particularly attractive for sharing executable code, and for documenting analysis extensively for reproducibility.

### Data Format

MathIOmica uses standard Mathematica expressions. In addition we created a simple structured data format termed an OmicsObject, Figure 2. This OmicsObject input format was created to facilitate data organization, with an eye for multiple datasets with different information included, such as identifiers to samples, identifiers to entities in the sample (e.g. gene names), measurements for the entities (e.g. intensities or gene expression), and any metadata the user may wish to keep or use in their analysis. The main format is two levels of associations, with outer keys matching sample labels, inner keys matching identifiers for the components, and values for each inner key taking two lists, one for measurements and another for metadata, as shown in Figure 2. The OmicsObject was designed to utilize 10.3+ Wolfram Language improvements to use associations (cf. dictionaries in Python). This data format allows rapid sample identification, measurement and metadata all maintained at various stages for user accessibility. MathIOmica includes specialized functions designed specifically for the OmicsObject data format to assist the user in controlling all aspects of the input data, Figure 1. Furthermore, an OmicsObject is an association of associations, and so the inbuilt powerful Wolfram Language Query function can be used directly to access and manipulate components.

### Data Processing

#### OmicsObject creation and manipulation

MathIOmica offers utilities to process mapped data and import them into an OmicsObject, Figure 1, including a graphical interface, Figure 4. The data format for the graphical interface importer can be any text delimited file, including files with comma separated values (csv), or tab delimited (tsv), as well as Excel spreadsheets (Microsoft), which are typical standard outputs in many computational applications or informatics software. Once the data has been imported or cast into an OmicsObject then the data may be processed as fit for each omics. This includes transformations such as quantile normalization, Box-Cox power transformations^43^, and filtering based on any field. Additionally the data can be tagged for low or missing values and filtered. For more customizable options the Applier function allows the user to apply any function of interest across the OmicsObject components. The workflow for the example multi-omics implementation in the documentation’s tutorials is shown in Figure 3 - we are also providing a printout version in Supplementary Note 1.

#### Time Series

- *Time Series from an OmicsObject*. From an OmicsObject simple operations can create a time series for each component, e.g. each gene. MathIOmica provides functions to facilitate the process, such as CreateTimeSeries and TimeExtractor. The functions assume an OmicsObject as an input for which times have been used as the sample labels strings (outer keys). The time series can be unevenly sampled and contain missing values as well.
- *Spectral Analysis*. Time series spectral analysis for missing data has been implemented through various standard approaches in MathIOmica. The main functionality exists to use a Lomb-Scargle transformation to handle uneven sampling and/or missing data, an approach that has been adapted from astronomy and used in the analysis and classification of dynamics in biological systems^17, 44–56^. Two main options are provided, the LombScargle function for reconstructing a periodogram directly, and additionally the Autocorrelation function for obtaining the autocorrelations through an inverse Fourier transform of the power spectrum for the data.
- *Classification and Clustering.* For sets of time series measurements, (e.g. gene expression levels, protein intensities, compound concentrations, temperature) we would like to identify groups of entities whose temporal behavior is similar. If we can classify temporal signals that show similar behavior into classes, then we can also look for associations/connections between the members of each class. The behavior of a given signal can be considered in terms of time or frequency (if we can Fourier transform the signal). The structure of the signal can then be described through autocorrelations, or equivalently its power spectrum (periodogram). Additionally, a signal can also be modeled with a time series model that has certain structure, e.g. autoregressive (AR), moving-average (MA), autoregressive moving-average (ARMA) etc. In this version, MathIOmica provides five different methods for time series classification through its TimeSeriesClassification function: Three methods use statistical cutoffs generated typically through simulation and are provided by the user: (i) Classification based on a Lomb-Scargle periodogram, classifying data into classes for time-series showing the same dominant frequency in their spectra; Classification based on autocorrelation, either computed by (ii) an inverse Fourier transform on the Lomb-Scargle periodogram to address missing data/uneven sampling or (iii) autocorrelation using interpolation (by default cubic) to address missing data/uneven sampling if any. Additionally two methods classify time series data into appropriate models, either (iv) by model kind, or (v) into classes that correspond to same model and parameters for the model. Following the classification, each class of time series can be clustered using hierarchical clustering using either TimeSeriesClusters or TimeSeriesSingleClusters. We should note here a subtlety when performing clustering of data series that are based on absolute values of intensity, (e.g. from a power spectrum/periodogram). In such cases, the sense (sign/phase) of the original time series cannot be detected directly from the intensity values. Therefore, such data vectors cannot distinguish correlated data from anti-correlated data that have the same periodogram structure. MathIOmica provides functionality to address this, through performing two tiers of clustering in TimeSeriesClusters, which can potentially distinguish the sense of each series, first clustering data using the periodograms to compute distances between components, and performing a secondary clustering on the results, using the original (non spectral) data for each time series that can assess the directionality of the data in real space.

#### Matrix Data Clustering

Standard clustering of data matrices can be performed by the function MatrixClusters in both the horizontal and vertical directions to identify groups based on similarities between the input series rows and columns.

### Annotation and Enumeration

MathIOmica provides annotations and over-representation analysis for gene ontology (GO) and KEGG pathways: (i) For GO analysis, the GOAnalysis function uses annotations (default is for human data) obtained from the Gene Ontology consortium. The annotation by default uses human data annotated with UniProt^57^ identifiers/accessions. Advanced function options also allow the user to use or directly download data for other species (examples with mouse and arabidopsis data are included in the documentation). An internal dictionary function can convert Gene Symbol and other identifiers obtained from the UCSC Browser^38^ tables to UniProt. (ii) In terms of KEGG pathway analysis, MathIOmica provides the KEGGAnalysis function. This uses annotations (default is for human data obtained from KEGG and by default uses human data annotated with KEGG Gene IDs(advanced options can also be used to utilize data from other species). Again, an internal dictionary function can convert Gene Symbol and UniProt identifiers to KEGG Gene IDs. Additionally, a molecular analysis is implemented for querying compounds against KEGG maps for metabolomics considerations.

Both GOAnalysis and KEGGAnalysis functions perform an over-representation (ORA) analysis, providing a p-value assessed by a hypergeometric function for membership in term categories/pathways. Additionally, a false discovery rate (FDR) cutoff for reporting is implemented, where adjusted p-values are computed (q-values) by a Benjamini Hochberg method^58^. The method and cutoffs can be customized (e.g. Bonferroni) by advanced users.

As mentioned above, MathIOmica includes a gene dictionary translation function, GeneTranslation, to convert identifiers between multiple databases. Additionally dictionaries are generated to provide GO term and KEGG pathway term descriptions.

### Visualization

- *Dendrograms and Heatmaps*. MathIOmica provides dendrogram/heatmap representations of clustering results. Separate functions are available to implement visualization for time series (with two-tier clustering [TimeSeriesDendrogramHeatmap] and single-tier clustering options [TimeSeriesSingleDendrogramHeatmap] available), as well as matrices [MatrixDendrogramHeatmap] that have been clustered. If classifications have been carried out, all classification output and graphs can be output simultaneously, e.g. by TimeSeriesDendrogramsHeatmaps (see Figure 4 and the inbuilt documentation).
- *KEGG Pathways*. A visual representation for KEGG pathways is implemented through use internally of the KEGG API. The returned data output can be a URL pointing to the online version of the pathway, that may be used in a browser, a downloaded figure, or a sequence of figures that may be used to generate animations in the case of time series data. Specific genes/proteins can be highlighted in the pathways, as well as represented in terms of intensities.
- *Mass Spectrometry Spectra.* MathIOmica provides a mass spectrum viewer for viewing .mzXML or .mzML^59, 60^ raw data. The viewer provides MS*^n^* viewing capabilities, filtering searches based on mass to charge ratios as well as retention times, viewing of precursor spectra and additionally summarizes the file metadata.

### Documentation and Examples

One distinct feature of MathIOmica is the utilization of Mathematica inbuilt system for documentation. MathIOmica was compiled so that its documentation is directly available in the Mathematica help system. This includes autocompletion in the system, templates for function use and direct access to definitions while developing code. Every function has at least a working example that can be evaluated in place, and documentation of all option choices for a given function. In all, more than 1000 pages of documentation and output are available. A printout of the function manual and tutorial are also available online at the MathIOmica websites.

#### iPOP examples

Data from the first integrative Omics Profiling^16, 17^ (iPOP) are used for inbuilt examples, comprised of dynamics from proteomics transcriptomics and metabolomics. Briefly, the data corresponds to a time series analysis of omics from blood components from a single individual. Different samples (from 7 to 21 included here) were obtained at different time points. The time points included here correspond to days ranging from 186th to the 400th day of the study. On day 289 the subject of the study had a respiratory syncytial virus (RSV) infection. Additionally, after day 301, the subject displayed high glucose levels and was eventually diagnosed with type 2 diabetes. The analyzed mapped data are used in these examples for illustrative purposes - these and additional dynamic omics data that will become available can also be accessed on MathIOmica’s main website. Various analyzed portions of the data are used in the documentation examples.

Furthermore, two full simple analyses are presented for the integrated full omics and streamlined transcriptome analyses as documentation tutorials. The main steps are depicted in Figure 3, and the tutorials are included as Supplemental Notes 1 (multiple omics) and 2 (transcriptome). Briefly, as shown in Figure 3, this simplified analysis example shows how each omic component can be first analyzed according to its own considerations towards a normal distribution and a time series of normalized intensities, relative to a healthy reference point. For each omic dataset a Lomb-Scargle periodogram classification identifies dominant frequency classes addressing periodicity, as well as Spike Max and Spike Min classes that correspond to singular events of high or low intensity. Cutoffs for the classification are provided from a bootstrap distribution construction (sampling with replacement) to compare against simulated randomized measurements. Following classification, data is clustered directly based on their periodograms (agglomerative clustering). Groups and subgroups are identified, with automatic group identification performed through a standard silhouette algorithm. The groups and subgroups in the different pattern categories are then checked for biological significance, looking for enrichment in GO categories and KEGG pathways. KEGG pathways are finally visualized and can be color coded to create animations corresponding to the intensities at different time points.

## Discussion

We have created MathIOmica, one of the first open source packages in the Wolfram Language to perform bioinformatics analyses. MathIOmica provides a general framework that is extensible to enable current and future workflows for integrating multiple data. This includes analysis of data, classification, annotation and visualization modules. The extensive inbuilt documentation provides an interactive environment for users to learn function usage and implementation.

We must note that the current version of MathIOmica is based entirely on established algorithms, which are also already available and implemented in other platforms/languages used widely by practicing bioinformaticians, especially through Bioconductor^22^. MathIOmica’s features are directed at an interdisciplinary audience, in addition to the practicing bioinformaticians, to include particularly mathematicians and physicists that use Mathematica in their research. It utilizes the high-level Wolfram language that is very familiar to quantitative scientists, and we believe this will facilitate novel development and introduce new algorithms into bioinformatics. Mathematica interfaces directly with R, Java and other languages, including the ability to work with C code, so it supplements other platforms and tools. The notebook interface is extremely attractive for prototyping, developing and sharing results, as well as a teaching platform.

We anticipate the next versions of MathIOmica to include further direct analysis of raw data, in addition to more advanced downstream classification and network analysis, as well as graphical user interfaces. We also plan to include a genome visualization, to locate differentially expressed genes in the genome, and merge structural and functional genomics in one package. All future versions and improvements will be provided for free at mathiomica.org and github repositories.

## Methods

### Code Base

MathIOmica was written exclusively in the Wolfram language. The package was built using Wolfram Workbench plugin for Eclipse (Mars) and built on Mathematica 10.4. It is also compatible with the newly released Mathematica 11. All source code is freely available and openly provided under an MIT open source license.

### Availability

MathIOmica is available for download as a Mathematica package at:

- https://mathiomica.org
- https://github.com/gmiaslab/mathiomica

The package contains the full open source code and documentation, which is released under an MIT open source license. All example files and data used therein are included as part of the package.

### Tutorial Supplementary Notes

MathIOmica’s documentation tutorials for multiple omics and transcriptome are provided online as Supplementary Notes 1 and 2 respectively.

## Acknowledgements

G.I.M. and research reported in this publication are supported by grants from Michigan State University and the National Human Genome Research Institute of the National Institutes of Health under Award Number 4R00HG007065. The content is solely the responsibility of the authors and does not necessarily represent the official views of the National Institutes of Health. R.R. is supported by a Paul and Daisy Soros Fellowship for New Americans. L.R.K.B. is supported by a Michigan State University AAGA Fellowship and a Michigan State University Enrichment Fellowship.

## Author contributions statement

G.I.M. conceived of and oversaw the study, carried out project planning, software design, tested and developed code, and wrote the manuscript. T.Y. was involved in computational work, wrote code and documentation, R.R. wrote code, L.R.K.B. wrote code and analyzed data, V.V.S. provided experimental data, C.C. designed and implemented the project website. G.I.M. is the senior author on this work. All authors reviewed the manuscript.

## References

1. Patti, G. J., Yanes, O. & Siuzdak, G. Innovation: Metabolomics: the apogee of the omics trilogy. Nat. Rev. Mol. Cell. Biol. 13, 263–269 (2012).

2. Mardis, E. R. Next-generation sequencing platforms. Annu. Rev. Anal. Chem. 6, 287–303 (2013).

3. Reuter, J. A., Spacek, D. V. & Snyder, M. P. High-throughput sequencing technologies. Mol. Cell 58, 586–597 (2015).

4. Wilhelm, M. et al. Mass-spectrometry-based draft of the human proteome. Nature 509, 582–587 (2014).

5. Kim, M.-S. et al. A draft map of the human proteome. Nature 509, 575–581 (2014).

6. Goodwin, S., McPherson, J. D. & McCombie, W. R. Coming of age: ten years of next-generation sequencing technologies. Nat. Rev. Gen. 17, 333–351 (2016).

7. ENCODE Project Consortium et al. An integrated encyclopedia of DNA elements in the human genome. Nature 489, 57–74 (2012).

8. 1000 Genomes Project Consortium et al. A global reference for human genetic variation. Nature 526, 68–74 (2015).

9. UK10K Consortium et al. The UK10K project identifies rare variants in health and disease. Nature 526, 82–90 (2015).

10. Collins, F. S. & Varmus, H. A new initiative on precision medicine. N Engl J Med 372, 793–5 (2015).

11. Dewey, F. E. et al. Phased whole-genome genetic risk in a family quartet using a major allele reference sequence. PLoS Genet 7, e1002280 (2011).

12. Jones, B. Genomics: personal genome project. Nature Publishing Group 13, 599 (2012).

13. Lesko, L. J. & Schmidt, S. Individualization of drug therapy: history, present state, and opportunities for the future. Clin. Pharmacol. Ther. 92, 458–66 (2012).

14. Whirl-Carrillo, M. et al. Pharmacogenomics knowledge for personalized medicine. Clin. Pharmacol. Ther. 92, 414–417 (2012).

15. McDonagh, E. M., Whirl-Carrillo, M., Garten, Y., Altman, R. B. & Klein, T. E. From pharmacogenomic knowledge acquisition to clinical applications: the PharmGKB as a clinical pharmacogenomic biomarker resource. Biomarkers in medicine 5, 795–806 (2011).

16. Mias, G. I. & Snyder, M. Personal genomes, quantitative dynamic omics and personalized medicine. Quant. Biol. 1, 71–90 (2013).

17. Chen, R. et al. Personal omics profiling reveals dynamic molecular and medical phenotypes. Cell 148, 1293–1307 (2012).

18. Mias, G. I. & Snyder, M. Multimodal Dynamic Profiling of Healthy and Diseased States for Future Personalized Health Care. Clin. Pharmacol. Ther. 93, 29–32 (2012).

19. Ghosh, S., Matsuoka, Y., Asai, Y., Hsin, K.-Y. & Kitano, H. Software for systems biology: from tools to integrated platforms. Nature Publishing Group 12, 821–832 (2011).

20. Moreau, Y. & Tranchevent, L.-C. Computational tools for prioritizing candidate genes: boosting disease gene discovery. Nat. Rev. Gen. 13, 523–536 (2012).

21. Hackl, H., Charoentong, P., Finotello, F. & Trajanoski, Z. Computational genomics tools for dissecting tumour-immune cell interactions. Nat. Rev. Gen. 17, 441–458 (2016).

22. Gentleman, R. C. et al. Bioconductor: open software development for computational biology and bioinformatics. Genome Biol. 5, R80 (2004).

23. Huber, W. et al. Orchestrating high-throughput genomic analysis with Bioconductor. Nat. Meth. 12, 115–121 (2015).

24. Cock, P. J. A. et al. Biopython: freely available Python tools for computational molecular biology and bioinformatics. Bioinformatics 25, 1422–1423 (2009).

25. Goecks, J., Nekrutenko, A., Taylor, J. & Galaxy Team. Galaxy: a comprehensive approach for supporting accessible, reproducible, and transparent computational research in the life sciences. Genome Biol. 11, R86 (2010).

26. Reich, M. et al. GenePattern 2.0. Nat. Genet. 38, 500–501 (2006).

27. Huang, D. W., Sherman, B. T. & Lempicki, R. A. Systematic and integrative analysis of large gene lists using DAVID bioinformatics resources. Nat Protoc 4, 44–57 (2009).

28. Shannon, P. et al. Cytoscape: a software environment for integrated models of biomolecular interaction networks. Genome Res. 13, 2498–504 (2003).

29. R Core Team. R: A Language and Environment for Statistical Computing (Vienna, Austria, 2013).

30. Wolfram Research, Inc. Mathematica, Version 10.4 (Wolfram Research, Inc., Champaign Illinois, 2015).

31. Wolfram, S. An Elementary Introduction to the Wolfram Language (Wolfram Media Inc, 2015).

32. Shapiro, B. E., Hucka, M., Finney, A. & Doyle, J. MathSBML: a package for manipulating SBML-based biological models. Bioinformatics 20, 2829–2831 (2004).

33. Baran, R. et al. MathDAMP: a package for differential analysis of metabolite profiles. BMC Bioinform. 7, 530 (2006).

34. Vilar, J. M. G. & Saiz, L. CplexA: a Mathematica package to study macromolecular-assembly control of gene expression. Bioinformatics 26, 2060–2061 (2010).

35. Allen, T. Detecting Differential Gene Expression Using Affymetrix Microarrays. Math. J. 15 (2013).

36. Hütt, M.-T. & Dehnert, M. Methoden der Bioinformatik. Eine Einf¨hrung zur Anwendung in Biologie und Medizin (Springer-Verlag, Berlin, Heidelberg, 2015).

37. Karolchik, D., Hinrichs, A. S. & Kent, W. J. The UCSC Genome Browser. Curr. Protoc. Bioinformatics Chapter 1, Unit1 4 (2012).

38. Speir, M. L. et al. The UCSC Genome Browser database: 2016 update. Nucleic Acids Res. 44, D717–25 (2016).

39. Ashburner, M. et al. Gene ontology: tool for the unification of biology. The Gene Ontology Consortium. Nat. Genet. 25, 25–29 (2000).

40. Gene Ontology Consortium. Gene Ontology Consortium: going forward. Nucleic Acids Res. 43, D1049–56 (2015).

41. Kanehisa, M. & Goto, S. KEGG: kyoto encyclopedia of genes and genomes. Nucleic Acids Res. 28, 27–30 (2000).

42. Kanehisa, M., Sato, Y., Kawashima, M., Furumichi, M. & Tanabe, M. KEGG as a reference resource for gene and protein annotation. Nucleic Acids Res. 44, D457–62 (2016).

43. Box, G. E. & Cox, D. R. An analysis of transformations. J. R. Stat. Soc. Series B Stat. Methodol. 26, 211–252 (1964).

44. Lomb, N. R. Least-squares frequency analysis of unequally spaced data. Astrophys. Space Sci. 39, 447–462 (1976).

45. Scargle, J. D. Studies in astronomical time series analysis. I - Modeling random processes in the time domain. Astrophys. J., Suppl. Ser. 45, 1 (1981).

46. Scargle, J. D. Studies in astronomical time series analysis. II-Statistical aspects of spectral analysis of unevenly spaced data. Astrophys. J. 263, 835–853 (1982).

47. Scargle, J. D. Studies in astronomical time series analysis. III-Fourier transforms, autocorrelation functions, and cross-correlation functions of unevenly spaced data. Astrophys. J. 343, 874–887 (1989).

48. Schimmel, M. Emphasizing difficulties in the detection of rhythms with Lomb-Scargle periodograms. Biol. Rhythm Res. 32, 341–5 (2001).

49. Van Dongen, H. P., Ruf, T., Olofsen, E., VanHartevelt, J. H. & Kruyt, E. W. Analysis of problematic time series with the Lomb-Scargle Method, a reply to ’emphasizing difficulties in the detection of rhythms with Lomb-Scargle periodograms’. Biol. Rhythm Res. 32, 347–54 (2001).

50. Bretthorst, G. L. Frequency Estimation and Generalized Lomb-Scargle Periodograms. In Statistical Challenges in Astronomy, 309–329 (Springer New York, New York, 2003).

51. Glynn, E. F., Chen, J. & Mushegian, A. R. Detecting periodic patterns in unevenly spaced gene expression time series using Lomb-Scargle periodograms. Bioinformatics 22, 310–6 (2006).

52. Caiado, J., Crato, N. & Peña, D. A periodogram-based metric for time series classification. Comput. Stat. Data Anal. 50, 2668–2684 (2006).

53. Zhao, W., Agyepong, K., Serpedin, E. & Dougherty, E. R. Detecting Periodic Genes from Irregularly Sampled Gene Expressions: A Comparison Study. EURASIP J. Bioinform. Syst. Biol. 2008, 1–8 (2008).

54. Gregory, P. C. (Philip Christopher), 1941. Bayesian logical data analysis for the physical sciences : a comparative approach with Mathematica support (Cambridge ; New York : Cambridge University Press, 2010).

55. Marcobal, A. et al. Metabolome progression during early gut microbial colonization of gnotobiotic mice. Sci. Rep. 5, 11589 (2015).

56. Wu, G., Anafi, R. C., Hughes, M. E., Kornacker, K. & Hogenesch, J. B. MetaCycle: an integrated R package to evaluate periodicity in large scale data. Bioinformatics btw405 (2016).

57. The UniProt Consortium. UniProt: a hub for protein information. Nucleic Acids Res. 43, D204–12 (2015).

58. Benjamini, Y. & Hochberg, Y. Controlling the false discovery rate: a practical and powerful approach to multiple testing. J. R. Stat. Soc. Series B Stat. Methodol. (1995).

59. Deutsch, E. mzML: a single, unifying data format for mass spectrometer output. Proteomics 8, 2776–7 (2008).

60. Martens, L. et al. mzML–a community standard for mass spectrometry data. Mol. Cell. Proteomics 10, R110 000133 (2011).

